# Resolving microbial microdiversity with high accuracy, full length 16S rRNA Illumina sequencing

**DOI:** 10.1101/010967

**Authors:** Catherine M. Burke, Aaron E. Darling

**Author notes:** Corresponding author: Catherine Burke. Corresponding author: Aaron Darling.

## Abstract

We describe a method for sequencing full-length 16S rRNA gene amplicons using the high throughput Illumina MiSeq platform. The resulting sequences have about 100-fold higher accuracy than standard Illumina reads and are chimera filtered using information from a single molecule dual tagging scheme that boosts the signal available for chimera detection. We demonstrate that the data provides fine scale phylogenetic resolution not available from Illumina amplicon methods targeting smaller variable regions of the 16S rRNA gene.

## INTRODUCTION

Amplifying and sequencing 16S rRNA genes from microbial communities has become a standard technique to survey and compare communities across space, time and environments. High-throughput sequencing methods have made bacterial community profiling routine and affordable, but at the expense of read length, with most platforms covering 250 to 600 bp of the ~1500 bp 16S rRNA gene. Depending on the region sequenced,these shorter fragments give variable taxonomic and phylogenetic resolution ^1^–^3^ and fails to resolve differences outside the targeted region, which may be ecologically relevant^4^–^6^.

Interpretation of 16S amplicon sequencing data is confounded by PCR and sequencing artefacts including chimeras, biased amplification and sequencing errors. Some of these artefacts can be removed computationally^7^, but nevertheless lead to errors that artificially inflate diversity estimates^8^–^10^ and mislead analysis.

## RESULTS

We present a method to sequence full-length 16S amplicons using Illumina technology. The technique incorporates randomized molecular tags on both ends of individual 16S template molecules prior to PCR amplification. Copies of the templates are fragmented and sequenced and the dual tag information is used to accurately re-assemble full length 16S gene sequences.In addition to generating longer sequences, this method offers large reductions in base calling errors,reduced amplification bias^11^ (see Supplementary materials), and a molecular signal supporting the identification and removal of artefacts generated by *in-vitro* recombination (chimeras).We evaluate this method by application to the skin microbiome and demonstrate improved phylogenetic resolution relative to short read sequencing of the V4 or the V1-4 regions. An overview of the method is shown in Figure 1.

**Figure 1.**
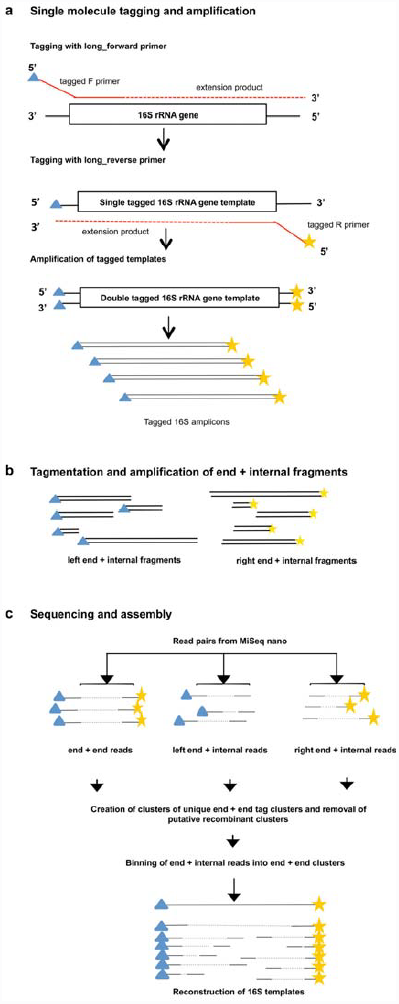
Overview of the Long-Read method. a) 16S rRNA gene template molecules are tagged with unique tags via two single rounds of annealing and extension with tagged forward and reverse primers (see Figure S1), that also contain Illumina adapter sequences. After removal of tagged primers, tagged templates are amplified via PCR using primers complementary to the adapter sequences. Libraries from one or more samples can then be pooled and sequenced on the MiSeq. Blue triangles and yellow stars indicate random tags of 10bp. b) Full-length 16S amplicon Illumina libraries are tagmented using the standard Nextera method, and two pools of products are amplified, which contain either the left end of the tagged amplicons and an internal region, or the right end of the amplicon and an internal region. This procedure adds Nextera adapters for sequencing at the internal end of the fragments. c) Both full length and tagmented libraries are paired end sequenced, and the unique molecular tags are used to computationally cluster sequences from the same progenitor 16S rRNA gene molecule for assembly of full length sequences.

Previous approaches for molecular tagging of the 16S rRNA gene have targeted individual variable regions, rather than sequencing of the whole gene, have incorporated tags at only one end of the amplicon^8^, or tags too short^10^ to permit the assembly and chimera filtering techniques proposed here. We tagged full-length individual 16S rRNA gene template molecules with unique molecular tags at both ends (Figure 1a) in DNA samples from foot skin swabs. This was achieved through primers incorporating a randomized 10 nucleotide sequence in addition to the gene priming region,sample specific barcodes, and adapters for Illumina sequencing (Figure S1). Tagged 16S rRNA gene templates were then amplified via PCR to give a mixed pool of 16S amplicons, where those molecules with the same combination of unique tags at either end were highly likely to have originated from the same template. The 10 base pair random sequence allows for 4^10^ ⍰ 1 million possible unique tags at each end of the tagged molecules, and ~ 1 trillion unique tag combinations.

After PCR amplification, aliquots of the 16S libraries were fragmented and prepared for Illumina sequencing using the standard tagmentation approach (Nextera XT, Illumina) ^12^. For the purpose of full length 16S sequencing, the PCR step was altered to selectively amplify only fragments containing one or the other of the original tagged 16S template molecule ends, coupled with a Nextera adapter introduced inside the original template via tagmentation. Sequencing of both the full length and the tagmented amplicon pools was then carried out on a single MiSeq Nano flow cell using standard primers, yielding 832,293 read pairs.

Paired end sequences from the full length amplicons, referred to as end + end sequences (n=42715),were used to determine which pairs of unique molecular tags from either end of the amplicons were from the same progenitor molecule. A clustering algorithm was used to define groups of frequently occurring molecular tags in the presence of sequencing error, resulting in 5085 clusters of end + end sequences.Putative *in vitro* recombination products were identified as low abundance clusters containing a molecular tag on one or both ends that was also observed in a higher abundance cluster (see online methods). Putative recombinant clusters were on average 29 times less abundant than the non-recombinants, and represented 4378 of the 42715 end + end sequences, indicating a recombination rate of 10.2%.These recombinant forms are a common problem in amplicon sequencing^13^, and are difficult to detect in the absence of molecular tags (as demonstrated on short read V4 sequences, see Supplementary results). Their removal ensures a more accurate representation of the true diversity captured from each sample.

Reads from the tagmented amplicon pools were then assigned to previously defined clusters of molecular tag pairs, and assemblies (see online methods) were performed on read clusters to yield consensus 16S sequences, ranging in length from 400 bp to 1378 bp (full length). In a single MiSeq Nano run of 832,293 read pairs, we assembled 2304 sequences, 70% of which were greater than 1300bp (Figure S5). Assuming linear scaling, the method would yield up to 80,000 full length 16S sequences on a 600 cycle MiSeq v3 kit, while a HiSeq 2500 might generate more than 480,000 full length 16S sequences in a single “rapid run” lane.

For comparison, the same samples were subject to sequencing of the V4 region (250 bp) of the 16S gene with an adaptation of a widely used protocol^14^ (see online methods), using a MiSeq Nano flow cell with 150 bp paired end reads. Figure 2 shows the results obtained using the V4 and Long-Read protocols.Higher average quality scores were obtained for the new sequencing protocol (68.6±17.2, PHRED scale, estimated by bcftools) as compared to the V4 sequencing (37.4±1.3).The 100-fold reduction in average error rate resulted from the increased depth of coverage of individual 16S molecules (Figure 2a-b).

**Figure 2.**
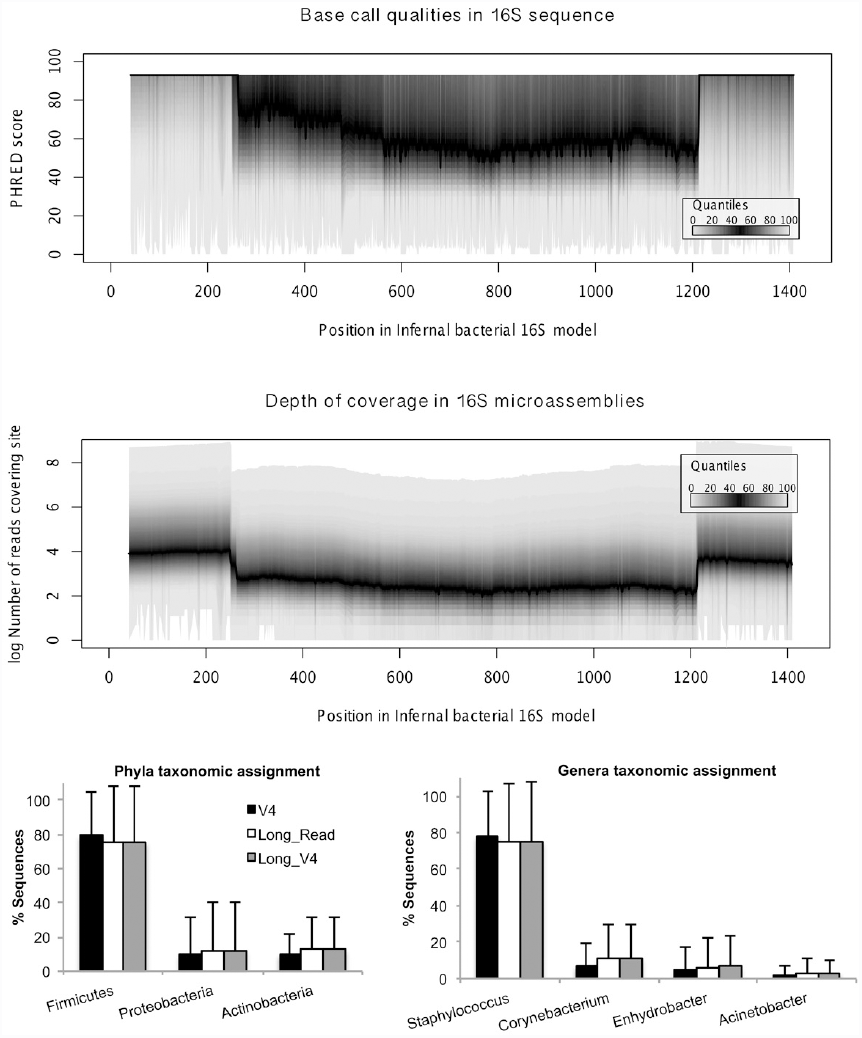
Results for Long-Read and V4 16S sequencing. a) Quality and b) coverage plots showing median PHRED quality scores and associated coverage across the length of the Long-Read 16S sequences.Higher qualities for the long-read sequences are associated with higher coverage,which is particularly apparent at each end of the amplicons (up to 200 and beyond 1200 bp), which were associated with one read from every read pair in the data set. c) Taxonomic assignments at the phylum and genus level of OTUs for V4, Long-Read and Long-V4 sequences (V4 region extracted from the Long-Read sequences). Similar taxonomic assignments are obtained across the V4 and Long-Read datasets. Only taxa comprising an average of more than 1% of sequences per sample are displayed. Bars represent the average value across 12 samples for each sequencing method, and error bars are the standard deviation.

Taxonomic assignments were compared between the V4 sequncing data and the V4 region of the full length data (longV4), to exclude differences in taxonomic assignment due to sequence length.Sequences were analysed using QIIME^15^ for OTU clustering and taxonomic assignment (see Supplementary methods). Similar taxonomic assignments were obtained at broad taxonomic classification levels (Phylum and Genus, Figure 2c) for both the V4 and long_V4_ sequences.

In order to evaluate how the new protocol influences phylogenetic resolution, we compared phylogenies constructed from sequences greater than 1300 bp to those constructed from the V4 (long_V4_) and V1-V4 (long_V1-4_) regions of the same data set. Phylogenies on full-length sequences
 were better resolved (2954 of a possible 3179 branches compared to 2686 for the V1-V4 region,and 2114 for the V4 region) with stronger support (Figure S5). While molecular tagging has previously been demonstrated to eliminate false microdiversity^8^,^10^, our data indicates it can also capture additional true microdiversity when applied to longer amplicons.

## CONCLUSION

We have demonstrated accurate reconstruction of full length 16S amplicon sequences from high throughput Illumina sequencing data. Although a protocol to sequence full length 16S amplicons on the Pacific Biosciences instrument has been demonstrated^16^, it can not at present match the throughput and cost efficiency of current Illumina platforms. We estimate nearly half a million full length 16S sequences could be generated per HiSeq 2500 lane.We note that the method of dual molecular tagging could be applied to any sequencing platform and any amplicon target to enhance chimera removal and reduce amplification bias and base calling error. This method, in conjunction with new algorithms^17^, will enable a finer understanding of population dynamics in microbial ecosystems.

## ACCESSIONS

Raw read data is available from SRA accession SRX655489. Assembled 16S sequences have been provided as a Supplementary data set, Long-Read.fasta

Software automating the process of sample demultiplexing, barcode clustering, and amplicon assembly is available from http://github.com/koadman/longas (to be made public).

## ACKNOWLEDGMENTS

We thank Josh Quick for advice on configuring the MiSeq to cluster long fragments, Paul Worden for assistance operating the MiSeq instrument, and Torsten Thomas for helpful discussions and suggestions during manuscript preparation.

